# Development and Validation of Fluorinated Amino Acid Parameters for use with the AMBER ff15ipq Protein Force Field

**DOI:** 10.1101/2022.01.06.475229

**Authors:** Darian T. Yang, Angela M. Gronenborn, Lillian T. Chong

## Abstract

We developed force field parameters for fluorinated aromatic amino acids enabling molecular dynamics (MD) simulations of fluorinated proteins. These parameters are tailored to the AMBER ff15ipq protein force field and enable the modeling of 4, 5, 6, and 7F-tryptophan, 3F- and 3,5F-tyrosine, and 4F- or 4-CF_3_-phenylalanine. The parameters include 181 unique atomic charges derived using the Implicitly Polarized Charge (IPolQ) scheme in the presence of SPC/E_b_ explicit water molecules and 9 unique bond, angle, or torsion terms. Our simulations of benchmark peptides and proteins maintain expected conformational propensities on the *μ*s-timescale. In addition, we have developed an open-source Python program to calculate fluorine relaxation rates from MD simulations. The extracted relaxation rates from protein simulations are in good agreement with experimental values determined by ^19^F NMR. Collectively, our results illustrate the power and robustness of the IPolQ lineage of force fields for modeling structure and dynamics of fluorine containing proteins at the atomic level.

## I. Introduction

While nuclear magnetic resonance (NMR) spectroscopy using the ^19^F nucleus is emerging as a powerful tool for measuring various structural and dynamical properties of fluorine-labeled biomolecules^1^, the availability of force fields for modeling the corresponding conformational dynamics at the atomic level has been limited. To address this unmet need, we have expanded upon the standard amino acids available in the AMBER ff15ipq protein force field^2^ and present a comprehensive set of parameters for the most commonly used fluorinated aromatic amino acids in ^19^F NMR experiments. Coupled with GPU-accelerated molecular dynamics (MD) simulations^3,4^, we can now begin to effectively integrate the results from ^19^F NMR experiments with atomistic MD simulations.

The fluorine nucleus has several features that render it a particularly useful NMR probe. Specifically, ^19^F is a 100% naturally abundant isotope with a high gyromagnetic ratio, which makes it almost as sensitive as ^1^H. Importantly, ^19^F chemical shifts span a very large range and are exquisitely responsive to the local chemical and electronic environment around the atom^5^. Fluorine is absent from virtually all naturally occurring biomolecules. Therefore, studies of fluorinated biopolymers can be carried out in any routine buffer system or environment without interference from background signals^6^. In general, only one or a handful of fluorine atoms are introduced into a biopolymer, overcoming the common handicap of spectral overlap in proton spectra that necessitated uniform labelling with ^13^C and ^15^N, in conjunction with 3- and 4D heteronuclear spectroscopy, for resonance assignments in bioNMR. These desirable properties have propelled ^19^F NMR based studies of biomolecular systems forward, such as investigations of thermodynamics^7^ and kinetics^8^, protein structure and dynamics^9–11^, or protein-protein and protein-ligand interactions^12–17^. Efforts are also ongoing towards ^19^F NMR method development, including relaxation optimization approaches^18,19^ to extend applicability to larger systems and to exploit ^19^F paramagnetic relaxation enhancement to determine long distances up to 35 Å^20^. ^19^F NMR is particularly useful in the pharmaceutical arena, guiding fragment-based screening^21^ and drug discovery/design^22^. Finally, a more recent and exciting direction is the movement of ^19^F NMR into a physiological environment, measuring spectra of fluorinated proteins directly in *Escherichia coli*^23^, oocyte^24^, and mammalian cells^25^.

Naturally, ^19^F NMR of proteins is not a panacea and has shortcomings and limitations. The 1.47 Å van der Waals radius of the fluorine atom lies between those of hydrogen (1.2 Å) and oxygen (1.52 Å). Therefore, fluorine is frequently substituted for hydrogens, hydroxyl groups, or carbonyl oxygens. While such substitutions are only weakly perturbing, with little to no effect on a protein’s structure and biological activity^7,10,11,26^, this is not necessarily the case and needs to be verified for each system under study^27^. Compared to more traditional NMR probes (^1^H, ^15^N, ^13^C), introducing a highly electronegative fluorine atom changes electronic surroundings and thereby local interactions, especially if the substituted H or OH was involved in hydrogen bonding. In addition, although resonance overlap is avoided, having only a few probes available, compared to thousands with traditional ^1^H, ^15^N, ^13^C labeling, limits the information content and several samples may need to be prepared where single fluorinated amino acids are judiciously placed to probe different parts of the biomolecule. MD simulations can help bridge this gap in information content at the all-atom level.

Here, we have developed new force field parameters for eight commonly used fluorinated aromatic amino acids, facilitating more accurate atomic-level simulations of ^19^F NMR observables. As indicated in **Figure 1**, our study focuses on 4, 5, 6, and 7 fluoro-tryptophan (W4F, W5F, W6F, W7F); 3, and 3, 5 fluoro-tyrosine (Y3F, YDF); as well as 4 fluoro- and 4 trifluoromethyl-phenylalanine (F4F, FTF). These fluorinated derivatives of tryptophan, tyrosine, and phenylalanine were selected because they are readily available and can be easily incorporated into proteins for ^19^F NMR^28–31^. Our parameters are intended for use with the AMBER ff15ipq force field and were derived using a general workflow that has been used previously for deriving new classes of non-canonical residues^32^. Consistent with the ff15ipq force field, we derived implicitly polarized atomic charges in the presence of explicit solvent. The implicit polarization of atomic charges has been demonstrated to be essential for modelling condensed-phase electrostatics in the absence of explicit polarization by fixed-charge force fields^2,33^, such as Drude oscillators^34^ and inducible multipoles^35^. Furthermore, application of the ff15ipq force field with the intended SPC/E_b_ water model yields more accurate rotational diffusion times of proteins, enabling direct calculation of NMR observables^2,36^. Using the GPU-accelerated AMBER MD engine^37,38^, we have extensively validated our force field parameters by simulating both peptides and proteins that include each of the eight fluorinated amino acids, yielding over 47 μs of aggregate simulation time. Our parameters maintain expected conformational propensities of both fluorinated peptide- and protein-based systems on the μs-timescale, and our relaxation rates from simulation agree with those from ^19^F NMR experiments.

**Figure 1.**
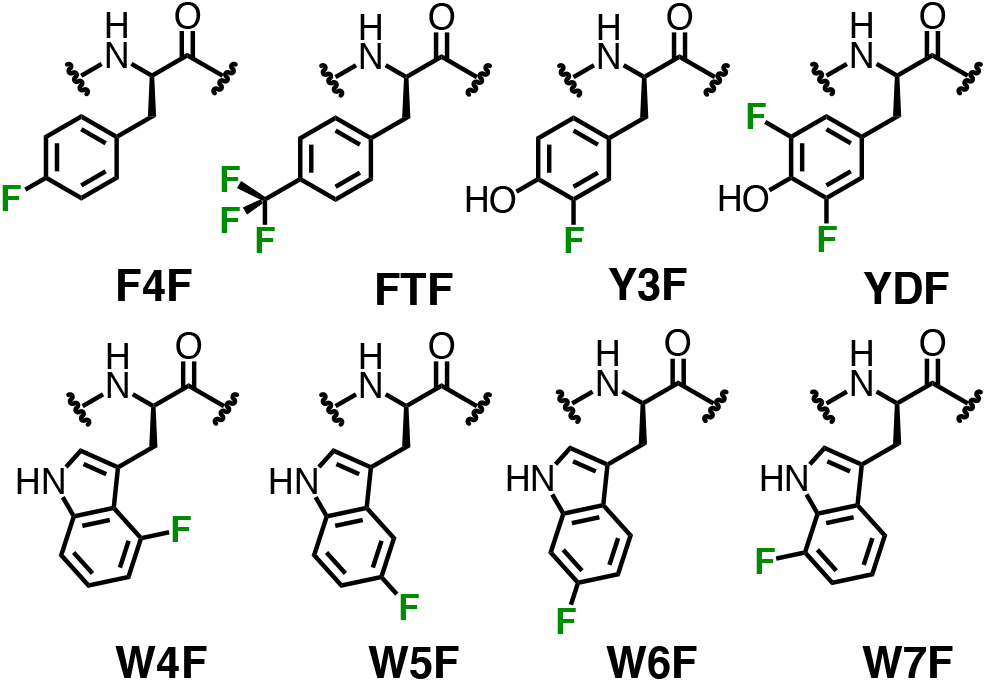
Eight commonly used fluorinated aromatic amino acids with their respective 3-letter identifier codes. The fluorine atom is highlighted in green.

Until now, parameters for the full set of fluorinated aromatic amino acids listed above for the AMBER ff15ipq protein force field have not been available, although recently, parameters for over 400 non-standard amino acids were added to the CHARMM36 protein and CHARMM general force fields^39^, including some for several fluorinated amino acids. The CHARMM parameters were derived by isolating only the sidechain atoms of the amino acid of interest and optimizing select water interactions based on quantum mechanical calculations at either the MP2/6-31G(d) or the HF/6-31G(d) level of theory. However, CHARMM parameters are not available for 7-fluoro-tryptophan and 4-fluoro-phenylalanine. In addition, parameters for apolar, non-aromatic, fluorinated amino acids are available for use with the AMBER ff14SB force field, which selectively optimized fluorine specific Lennard-Jones parameters and derived atomic charges at the RHF/6-31G* level of theory^40,41^. Also, for use with the ff14SB force field, a set of unnatural phenylalanine and tyrosine derivatives was parameterized which included 3, 5 fluorotyrosine^42^. Atomic charges for these parameters were based on electrostatic potential calculations of a dipeptide molecule at the HF/6-31G* level of theory. Other efforts to develop parameters for simulating fluorinated proteins have been primarily ad hoc and single-use cases^43–45^.

## II. The AMBER ff15ipq force field

Parameters for the eight different fluorinated amino acids were developed for use with the AMBER ff15ipq protein force field, the latest version of the Implicitly Polarized Charge (ipq) force field lineage.^2,46^ These parameters are also compatible with the ff15ipq-m force field for protein mimetics^32^. Ipq force fields feature implicitly polarized atomic charges, with each charge optimized to reproduce the mean field electron density of the molecule in explicit solvent^47^. Notably, the ff15ipq force field is parameterized using the three-point, explicit SPC/E_b_ (extended simple point charge) water model^48^, which reproduces the experimental rotational diffusion of proteins and can therefore yield reasonable dynamical observables such as those measured by NMR without the computational burden associated with four-point water model alternatives.

The original motivation behind the ff15ipq force field was to yield more accurate propensities for salt bridge formation, which is a common limitation of most contemporary fixed-charge force fields^33^. The final ff15ipq implementation was a complete rederivation of its predecessor^46^, including a greatly expanded torsion and angle parameter set, atomic charges derived at the MP2/cc-pVTZ level in explicit solvent, and adjustments to atomic radii for polar hydrogens. The ff15ipq force field reproduces the experimental probabilities of salt-bridge formation, while maintaining the secondary structure of stably folded and disordered systems on the μs-timescale, and faithfully predicts J-coupling constants for a penta-alanine peptide as well as NMR relaxation rates for protein systems^2^.

Additional developments involved a modification of methyl side chain rotational barriers that help to accurately predict methyl relaxation rates^49^, as well as an expanded force field (ff15ipq-m) for modeling four classes of artificial backbone units, including D- and Cα-methylated amino acids, β-amino acids and two cyclic β residues^32^. These developments are implemented in both the AMBER^37,38,50^ and open-source OpenMM^51^ GPU-accelerated biomolecular simulation software packages, and are accessible to other such packages through the ParmEd program^52^ for format conversion.

Our new parameters for the eight fluorinated aromatic amino acids were derived using a workflow designed to be consistent with the parent ff15ipq derivation process. As done for the ff15ipq-m force field, one minor difference pertained in the generation of dipeptide conformations for both the charge and bonded parameter derivation: to increase and safeguard conformational diversity, we progressively restrained the backbone torsions at evenly spaced intervals before energy-minimizing each restrained conformation^32^. In the parent ff15ipq force field, conformations were generated using high-temperature MD simulations^2^.

## III. Methods

### A. Derivation of IPolQ atomic charges

For each fluorinated residue, IPolQ atomic charges were derived using a four-step iterative procedure until convergence was reached (**Figure 2**):

1. **Generate a set of conformations.** For a blocked dipeptide (a single amino acid with capping groups) containing the fluorinated residue of interest with acetyl (Ace) and N-methylamide (Nme) N- and C-terminal capping groups, respectively, 20 conformations were generated by progressively restraining backbone Φ/Ψ torsion angles from −180° to 180° using a force constant of 32 kcal/mol. Only for the first iteration, the initial set of atomic charges were derived using the AM1-BCC charge method^53^. Each restrained conformation was then subjected to energy minimization and solvated using a truncated octahedral box of SPC/E_B_ water molecules with at least a 12 Å clearance between the solute and the edge of the box. After another round of energy minimization, each solvated system underwent a two-stage equilibration which included a 10 kcal/mol Å^2^ positional restraint on the fluorinated amino acid. In the first stage, 20 ps of dynamics were carried out at constant temperature (25°C) and volume. In the second stage, 100 ps of dynamics were carried out at constant temperature (25°C) and pressure (1 atm). A final 500 ps simulation was performed at constant temperature (25°C) and volume during which the solute remained fixed, and the solvent coordinates were used to generate a distribution of point charges that represent the solvent reaction field potential. This distribution consisted of an inner cloud of point charges based on the coordinates of the solvent molecules within 5 Å of the solute and three outer shells of point charges that reproduce contributions to the solvent reaction field potential from the periodic system beyond 5 Å.
2. **Calculate the electrostatic potential of each conformation in vacuum and explicit solvent**. Two sets of QM electrostatic potential calculations were carried out for each conformation of the fluorinated amino acid at the MP2/cc-pVTZ^54–57^ level of theory using the ORCA 4.2.0^58^ software package. The first set of QM calculations were in vacuum, and the other set included the solvent reaction field potential to represent the surrounding explicit solvent molecules.
3. **Fit the average electrostatic potential over all conformations to atomic charges, both in vacuum and explicit solvent**. All eight aromatic fluorinated amino acids were fit together using the fitq module of the mdgx program^50^, with atomic charges of the Ace and Nme capping groups fixed to net neutral values during the process.
4. **Average the vacuum-phase and solvent-phase point charges to obtain implicitly polarized atomic charges.** This averaged charge set was used as the starting point for the next iteration of charge generation and fitting as described above.

**Figure 2.**
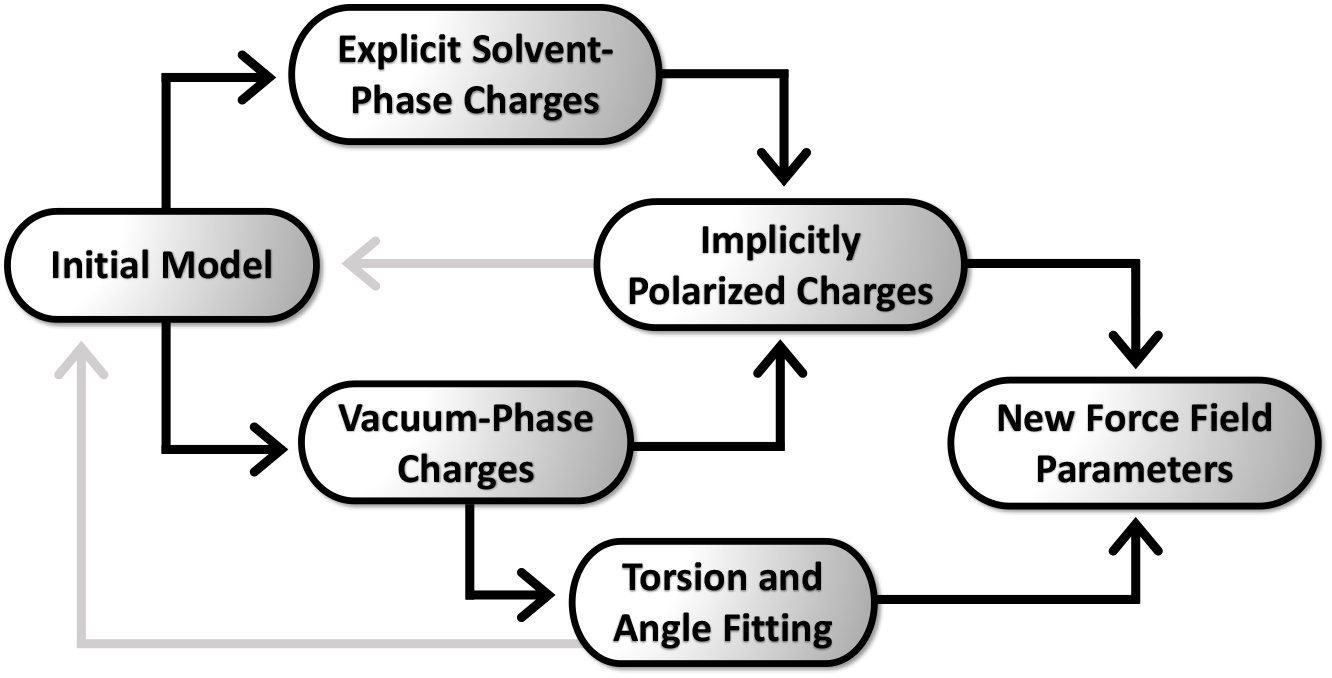
Workflow for developing ff15ipq force field parameters. Starting from a model consisting of multiple coordinate and topology files, the IPolQ charge derivation protocol is used to fit atomic charges from electrostatic potential calculations in both vacuum and explicit solvent phases. These charges are averaged, implicitly modeling solvent polarization effects. The vacuum-phase charges are used to fit bonded parameter terms to the vacuum-phase QM targets and used iteratively with the implicitly polarized charges, as indicated by the gray arrows, to generate a new initial model until the parameters are self-consistent.

The above procedure was repeated four times to reach convergence of the partial atomic charge values within 10% of the previous iteration’s values.

### B. Generation and fitting of the bonded parameter dataset

For each fluorinated amino acid, bonded parameters were derived using an iterative, multi-step procedure: (i) 1000 conformations of the blocked dipeptide were generated in vacuum, taking trial torsion and angle parameters from the parent ff15ipq force field, if available, or otherwise from the general amber force field 2^59^. Atomic charges were obtained from the vacuum phase set of converged point charges as described above. Each conformation was generated by progressively restraining backbone Φ and Ψ torsion angles of the fluorinated dipeptide between −180° and 180° using a force constant of 32 kcal/mol. After energy minimization Φ and Ψ distributions of the conformation set were plotted to ensure extensive sampling of the configurational space. (ii) For each conformation, the quantum mechanical (QM) single point energy was calculated at the MP2/cc-pVTZ level of theory, along with the molecular mechanical (MM) energy. Linear least-squares fitting for the entire set of bond, angle, and torsion parameters was run using mdgx^60^. (iii) Force field bonded parameters such as torsional barrier heights, angle equilibria, angle stiffness, bond length equilibria, and bond stiffness were adjusted to minimize the error between the QM and MM energies. Using the updated parameter set, a second round of conformer generation and QM energy calculations was carried out in the absence of restraints to prevent becoming trapped in local minima. This new set of conformations excluded redundant conformations, as defined by those with MM energies that differ by <0.01 kcal/mol. Another round of fitting was then performed using both sets of conformations and their respective energies to obtain the final parameter set for each iteration. Steps (i-iii) were repeated until the root-mean-square error (RMSE) between the QM and MM energies was less that 1% from the previous iteration (**Figure 2**).

The accuracy of the molecular mechanical energies (U_MM_) produced using the optimized parameters for each fluorinated amino acid was assessed by comparing them to the respective QM energies (U_QM_) for all generated conformations (**Figure 3**). During each iteration of the above fitting procedure, optimization of the bonded parameters was monitored by the RMSE, which reached final values of 1.08 - 1.86 kcal/mol for all eight residues. These errors are slightly higher, on average, than those of the non-fluorinated tryptophan, tyrosine, and phenylalanine counterparts from the canonical ff15ipq force field^2^, which may be due to fewer parameters in the fitting procedure and/or the presence of electronegative fluorine atom(s).

**Figure 3.**
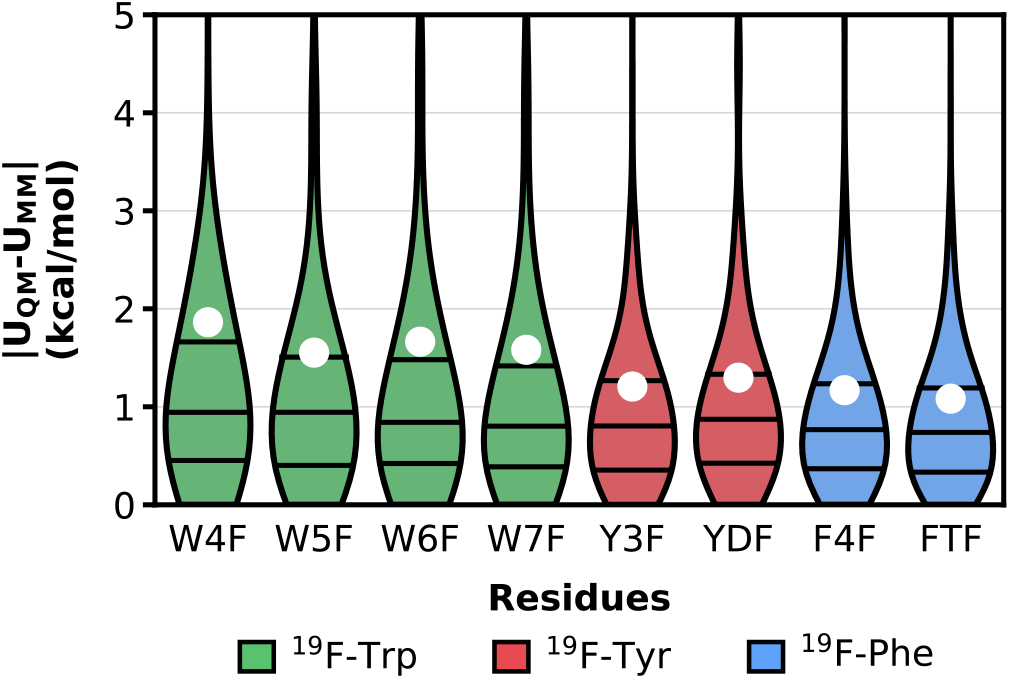
Distributions of residuals between quantum mechanical (U_QM_) energies and molecular mechanical (U_MM_) energies for the eight fluorinated Ace-Xaa-Nme dipeptides where Xaa is the fluorinated residue of interest. The 25^th^, 50^th^, and 75^th^ percentiles are represented by horizontal lines and root-mean-square-error values are represented by white filled circles. Each dataset is colored according to the original residue type indicated in the legend.

### C. Preparation of validation systems

Models of fluorinated and canonical tetrapeptides were built using Avogadro^61^ and tleap^60^. To generate the 4, 5, 6, and 7F-Trp modified protein systems, the atomic coordinates of cyclophilin A (CypA; Protein Data Bank^62^ (PDB) ID: 3K0N)^63^ were modified using Chimera^64^, with fluorine atoms substituted individually for hydrogens in Trp121 at the respectively numbered positions on the indole ring. Each system was solvated in a truncated octahedral box of explicit SPC/E_b_^48^ water molecules with a 10 Å clearance between the solute and the edge of the box for the peptides and a 12 Å clearance for the proteins. All systems with unpaired charges were neutralized by adding Na^+^ or Cl^-^ ions, treated with Joung and Cheatham ion parameters^65^. Protonation states for ionizable residues were adjusted to represent the major species present at pH 6.5 to match the experimental NMR conditions^66^.

### D. Umbrella sampling of fluorinated and canonical peptides

To validate the conformational preferences of each fluorinated residue, umbrella sampling simulations were carried out for a tetrapeptide (a tripeptide with capping groups (Ace-Ala-Xaa-Ala-NMe) containing a fluorinated amino acid at the central residue Xaa. Ramachandran plots were generated for each tetrapeptide by calculating the potential of mean force as a function of the Φ and Ψ torsion angles of the fluorinated residue. Prior to umbrella sampling, each tetrapeptide was solvated and equilibrated as described above for unrestrained simulations, differing only in the duration of the final equilibration stage, which was 100 ps instead of 1 ns. Each window was then subjected to a 200 ps incrementally restrained equilibration prior to a 2 ns restrained simulation at constant temperature (25 °C) and pressure (1 atm). The Φ and Ψ torsions of the central residue were restrained using a harmonic penalty function with a force constant of 8 kcal/mol rad^2^ for each window with 10° intervals about each torsion angle. This restraint scheme resulted in a series of 1296 windows for each set of two torsions, cumulating into 22.8 μs of aggregate simulation time for the eight fluorinated residue classes and 10,368 windows. From each set of 1296 windows, the unbiased potential of mean force was reconstructed using the weighted histogram analysis method (WHAM)^67–69^.

### E. MD Simulations of fluorinated and wild-type CypA

Simulations of the 4, 5, 6, 7F-Trp fluorinated and the wild-type CypA proteins were carried out using the GPU-accelerated pmemd module of the AMBER 18 software package^37,38,50^, ff15ipq force field^2^, and our new fluorinated amino-acid parameters. Each system was initially subjected to energy minimization followed by a three-stage equilibration. In the first stage, a 20 ps simulation was carried out at constant volume and temperature (25 °C) in the presence of solute heavy atom positional restraints using a harmonic potential with a force constant of 1 kcal/mol Å^2^. In the second stage, a 1 ns simulation was carried out at constant temperature (25 °C) and pressure (1 atm) using the same harmonic positional restraints. Finally, an unrestrained 1 ns simulation was carried out before performing a 1 μs production simulation with both constant temperature (25 °C) and pressure (1 atm). Five production simulations were run for each CypA protein, yielding 25 μs of aggregate simulation time.

Temperatures were maintained using a Langevin thermostat with a frictional constant of 1 ps^−1^, while pressure was maintained using a Monte Carlo barostat with 100 fs between system volume changes. Van der Waals and short-range electrostatic interactions were truncated at 10 Å, while long range electrostatic interactions were calculated using the particle mesh Ewald method^70^. To enable a 2 fs time step, all CH and NH bonds were constrained to their equilibrium values using the SHAKE algorithm^71^. Coordinates were saved every ps for analysis in CPPTRAJ^72^.

### F. Backbone conformations of fluorinated amino acids in the PDB

A “ligand expo” search^73^ was carried out to determine whether each fluorinated amino acid of interest was present in PDB deposited structures. The corresponding three-letter identifiers in the PDB were as follows: 4FW, FTR, FT6, F7W, PFF, 55I, YOF, F2Y; which are equivalent to the following identifiers in the current study: W4F, W5F, W6F, W7F, F4F, FTF, Y3F, and YDF (**Figure 1**), respectively. Each identifier was associated with at least one structure and was used to construct a query for structures that included the fluorinated residue as a part of a polymer chain, avoiding fluorinated ligand molecules. For structures determined by X-ray crystallography, only those with resolution ≤ 2.5 Å were included, resulting in a total of 56 structures with at least one fluorinated residue each. The backbone Φ and Ψ torsion angles were then calculated using a custom Python script, omitting the torsion angles belonging to fluorinated C-terminal and N-terminal residues.

### G. Calculation of ^19^F NMR relaxation rates

^19^F longitudinal (*R*_1_) and transverse relaxation rates (*R*_2_) were calculated from MD simulations for each of the four ^19^F-Trp CypA protein systems using equations 1-4^5,74^. Both types of relaxation rates are affected by dipole-dipole (DD) interactions and chemical shift anisotropy (CSA). The influence of DD interactions is described by equations 1 and 2:

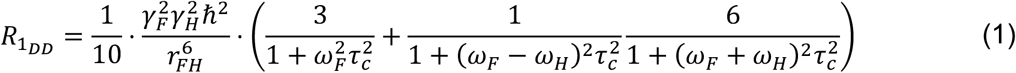

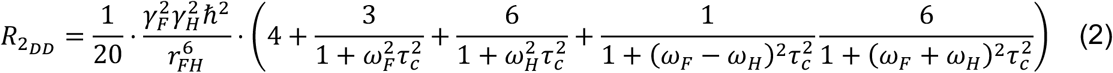

where *γ* is the gyromagnetic ratio of fluorine or hydrogen, 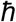 is the reduced Plank’s constant, *ω* is the Larmor frequency of fluorine or hydrogen, and *τ_c_* is the rotational correlation time.

For each frame in the MD simulations, the distances between each hydrogen and the fluorine atoms within a 3 Å radius around the fluorine were calculated (*r_FH_*) and used in equations 1 and 2 to calculate the relaxation rate contribution to fluorine from each nearby hydrogen. These contributions were summed to account for the influence of all surrounding hydrogen dipoles.

The influence of CSA effects is described by equations 3 and 4:

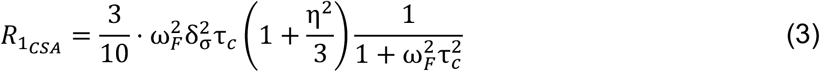

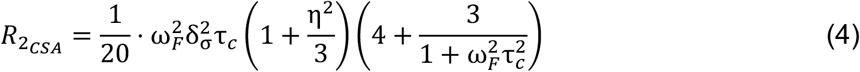

where *δ_σ_* is the reduced anisotropy and *η* is the asymmetry parameter, as described in Haeberlen^75^ convention by the following equations:

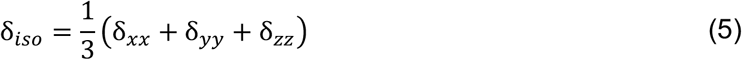

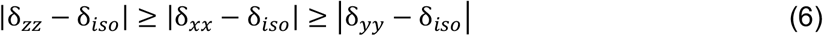

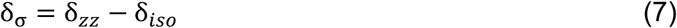

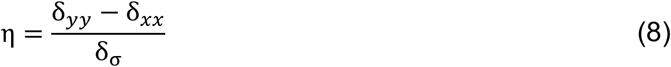

The final relaxation rate values were then calculated as the sum of the individual DD and CSA based components, where *R*_1_ = *R*_1_*DD*__ + *R*_1_*CSA*__ and *R*_2_ = *R*_2_*DD*__ + *R*_2_*CSA*__.

These equations assume the following approximations: (i) the protein tumbles isotropically with the rotational correlation time remaining constant and the fluorinated side chain motion is governed by the same overall protein rotational correlation; (ii) fluorine-hydrogen distances close the F atom remain constant and can be described by a single value; (iii) cross-correlation interactions^76^ are not accounted for, even though a 10-25% cross-term contribution to the total *R*_2_ relaxation rate may be possible^66^.

For all CypA variants a single rotational correlation time (*τ_c_*) of 8.2 ns^77^ was used, along with the ^19^F CSA values from solid-state magic angle spinning (MAS) NMR experiments for 4, 5, 6, or 7-fluoro-tryptophans^78^ (**Table S2**) to calculate *δ_σ_* and *η*. To sample a representative ensemble, only the last 800 ns of each trajectory, from each independent 1 μs production simulation, was used. All fluorine relaxation calculations were carried out using a custom-made Python program, which is open-source and freely available online (https://github.com/darianyang/fluorelax).

## IV. Results and Discussion

Our parameters for the eight fluorinated aromatic amino acids were derived using the IPolQ workflow (**Figure 2**). Briefly, a set of conformations for a capped peptide with the fluorinated residue of interest was generated, and electrostatic potentials of each conformation were calculated quantum mechanically, both in vacuum and in the presence of explicit solvent. The resulting set of vacuum and solvent phase atomic charges were then optimized to reproduce the respective electrostatic potential calculations, before being averaged to obtain an implicitly polarized charge set. Along with van der Waals interactions, these atomic charges estimate the non-bonded contributions of the force field. With the vacuum optimized charges, the bonded terms of the force field, such as bond angles and dihedrals, were fit to minimize the error between the molecular mechanical and the quantum mechanical energies of each conformation. Both the atomic partial charges and the bonded parameter derivation steps were repeated until they were self-consistent. To validate our force field parameters, we initially carried out peptide simulations to explore the changes in the conformational free energy landscape of each fluorinated residue. We then performed protein-based simulations of 4, 5, 6, and 7F-Trp substituted Cyclophilin A (CypA) and compared our simulation results with those of the native CypA protein. Finally, we calculated NMR relaxation rates from our MD simulation data of CypA and compared these rates to the respective, experimentally determined ^19^F relaxation rates from NMR.

### A. Conformational preferences of individual fluorinated residues

Backbone conformational preferences for the tetrapeptide systems (Ace-Ala-Xaa-Ala-Nme) were assessed using umbrella sampling simulations in which the central residue (Xaa) backbone dihedrals were progressively restrained (**Figure 4**). In umbrella sampling, a chosen reaction coordinate is initially divided into a series of windows or sections and a harmonic restraint is applied. In our case, this coordinate was the backbone Φ and Ψ dihedrals of the central residue. We choose 10° intervals from −180° to 180° using a force constant of 8 kcal/mol rad^2^ to ensure that the reaction coordinate remained within the center of the window during MD simulations (see methods section E). A series of histograms along the reaction coordinate was generated, and because these distributions were overlapping, our umbrella sampling parameters were determined to be exhaustive and allowed for the corresponding unbiased free energy landscape to be recovered using WHAM ^67–69^. Umbrella sampling was carried out for the fluorinated amino acid containing peptides as well as their non-fluorinated counterparts (Trp, Tyr, Phe), as depicted in the relative free energy difference plots (**Figure 4**). In addition, we compared our simulation results to the experimentally observed conformations for each fluorinated variant, extracted from 56 protein structures deposited in the PDB.

**Figure 4.**
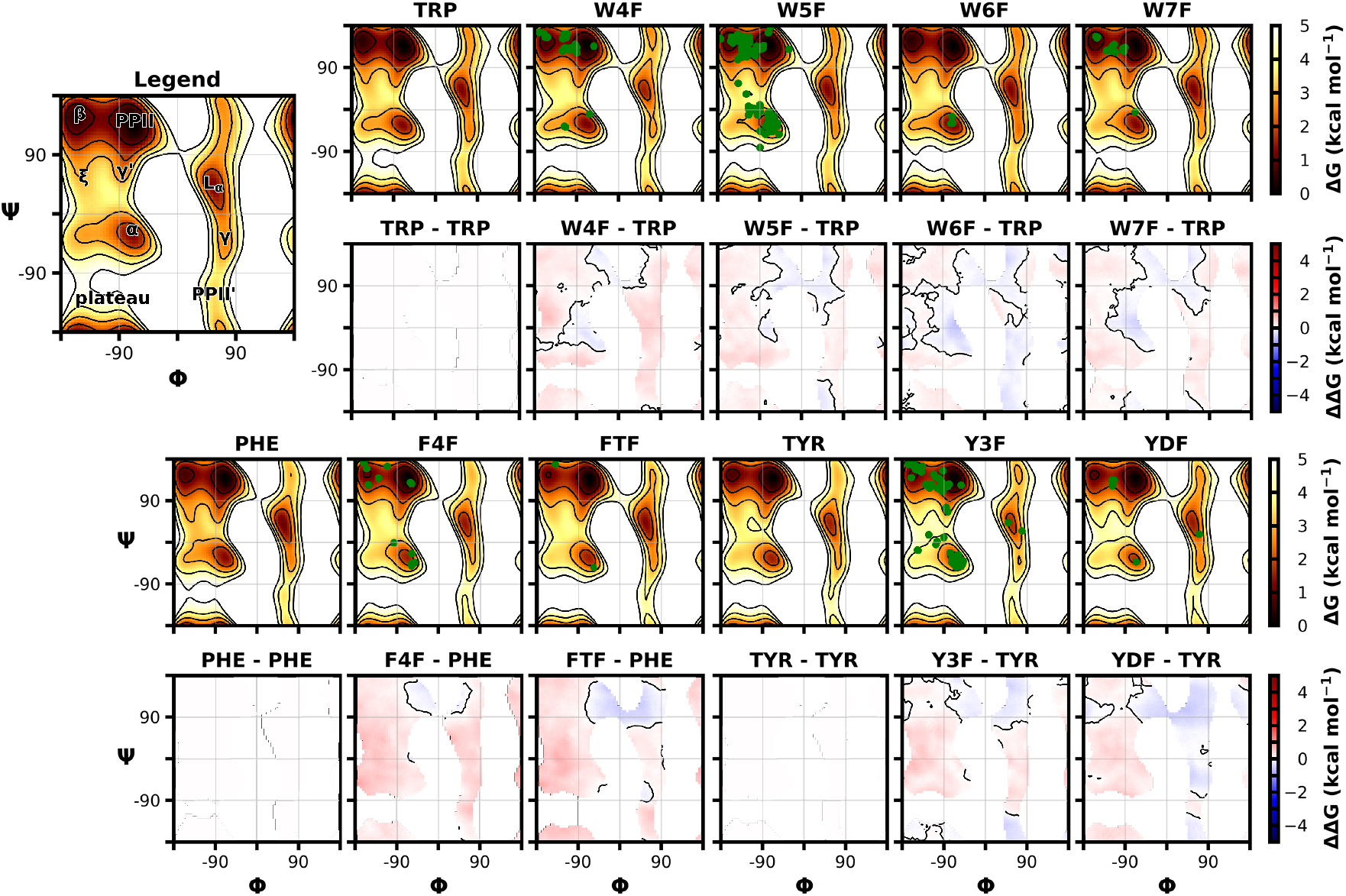
Free energy profiles for the central residue of tetrapeptides as a function of backbone Φ and Ψ torsions about the central residue, as well as the difference in free energy relative to the canonical non-fluorinated residue containing tetrapeptide. Each system is an Ace-Ala-Xaa-Ala-NMe peptide which underwent umbrella sampling simulations with subsequent application of WHAM. The green filled circles represent the backbone torsion angles observed for crystal structures that contain the same fluorinated residue as the tetrapeptide in their protein structure.

Our results show that most free energy barriers between different secondary structures were either unperturbed, or only slightly increased for the fluorinated residues relative to their non-fluorinated counterparts, indicating less favorable sampling of that region. An exception is the area between the poly-proline II (Φ ≈ −70°, Ψ ≈ 140°) and left-handed alpha helical (Φ ≈ 60°, Ψ ≈ 40°) regions, which were more favorably sampled in all cases for fluorinated residues. For tyrosine and phenylalanine variants, we saw this favorable sampling amplified with more fluorine atoms present in the rings, such as with tyrosine possessing one or two fluorine atoms and phenylalanine with a single fluorine or the trimethyl fluorine group. For tryptophan, our resulting energy landscapes produced a similar trend, except that the alpha helical (Φ ≈ −70°, Ψ ≈ −20°) and y’ (Φ ≈ −80°, Ψ ≈ 60°) regions were more favorably sampled, with fluorination at the six-position of the indole ring exhibiting the largest difference. All dihedral angles of fluorinated residues remained consistent with those experimentally observed in structures deposited in the PDB. The backbone Φ and Ψ torsion angle energies for fluorine containing peptides are very similar to non-fluorinated peptides, with ΔΔG values ranging from −0.63 to 0.64 kcal/mol for the tryptophan variants, −0.55 to 0.65 kcal/mol for the tyrosine variants, and −0.53 to 0.88 kcal/mol for the phenylalanine variants (**Table S2**). The average ΔΔG of our non-fluorinated versus fluorinated peptides are all close to zero, further supporting our conclusion that only minimal perturbations are induced upon fluorine substitution. The only exception is for 4-fluoro-phenylalanine, for which an average ΔΔG value of 0.273 ± 0.186 kcal/mol was observed. Overall, these findings are consistent with previous studies where fluorinated tryptophan^11,79^, tyrosine^80^, and phenylalanine^7^ substitutions introduced minimal to no differences in the global protein structure or the local dihedral angles around the fluorinated residue.

### B. Simulations of fluorinated Cyclophilin A

To evaluate our force field parameters in the context of a protein, we carried out multiple μs-timescale simulations for each of four variants of cyclophilin A (CypA), a 18.3 kDa peptidyl-prolyl isomerase that is a known host-factor for HIV-1 infection^81^. The variants are singly fluorinated at four different indole ring positions of Trp 121, which is close to the active site of the protein. These fluorinated variants of CypA have been studied previously^66^ and serve as useful benchmark proteins for our simulation studies.

Our data show that both wild-type and the fluorinated CypA variants all remained stable over the course of multiple μs-timescale simulations (**Figure 5**), and only small deviations are noted. For the wild-type CypA simulations, the average backbone RMSD value is 1.37 ± 0.37 Å (average ± one standard deviation), while the fluorinated Trp121 CypA variants exhibited average backbone RMSD values of 1.28 ± 0.23 Å (4F-Trp CypA), 1.22 ± 0.27 Å (5F-Trp CypA), 1.35 ± 0.31 Å (6F-Trp CypA), and 1.21 ± 0.20 Å, (7F-Trp CypA). RMSD values for each individual 1 μs simulation as a function of simulation time are shown in **Figure S2**.

**Figure 5.**
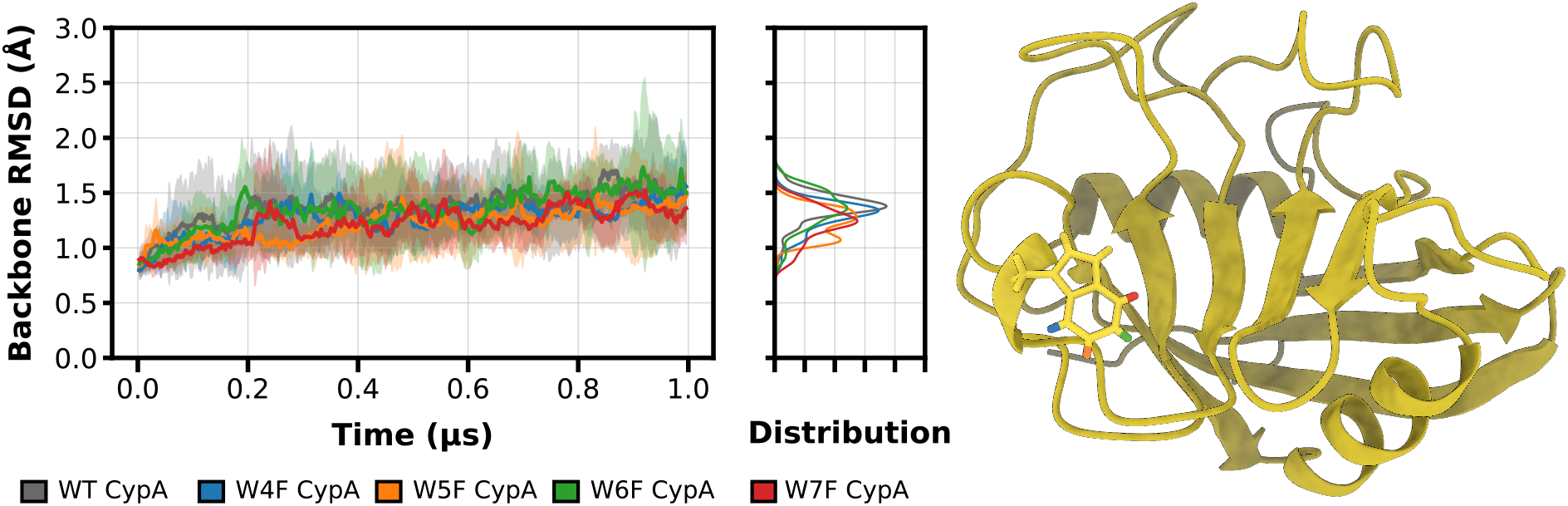
Average backbone RMSD values from the experimental X-ray structure (PDB: 3K0N)^63^ coordinates for native and fluorinated CypA proteins as a function of simulation time. Data shown is based on five independent 1 μs replicates, with one standard deviation of each time point shown in transparent coloration. The distribution of RMSD values is depicted in the middle panel and the protein structure being simulated is depicted on the right with individual fluorine positions on the TRP121 indole ring colored according to the legend.

In addition to backbone RMSD values, we also assessed the dihedral angle propensities of our aggregate simulation data by generating Ramachandran (Φ and Ψ) and Janin (χ_1_ and χ_2_) plots. All dihedral probabilities for the fluorinated CypA proteins remained within the same distribution as that of wild-type CypA (**Figure 6**). Only for 6F-Trp, a slightly larger range of conformational sampling around the alpha helical region in the Ramachandran plot and a slightly restricted conformational sampling distribution in the Janin plots was noted. These findings are similar to those with our peptide systems, where the 6F-Trp substitution also indicated a change in sampling of the free energy landscape near the alpha helical region, compared to the other fluorinated tryptophan moieties (**Figure 4**).

**Figure 6.**
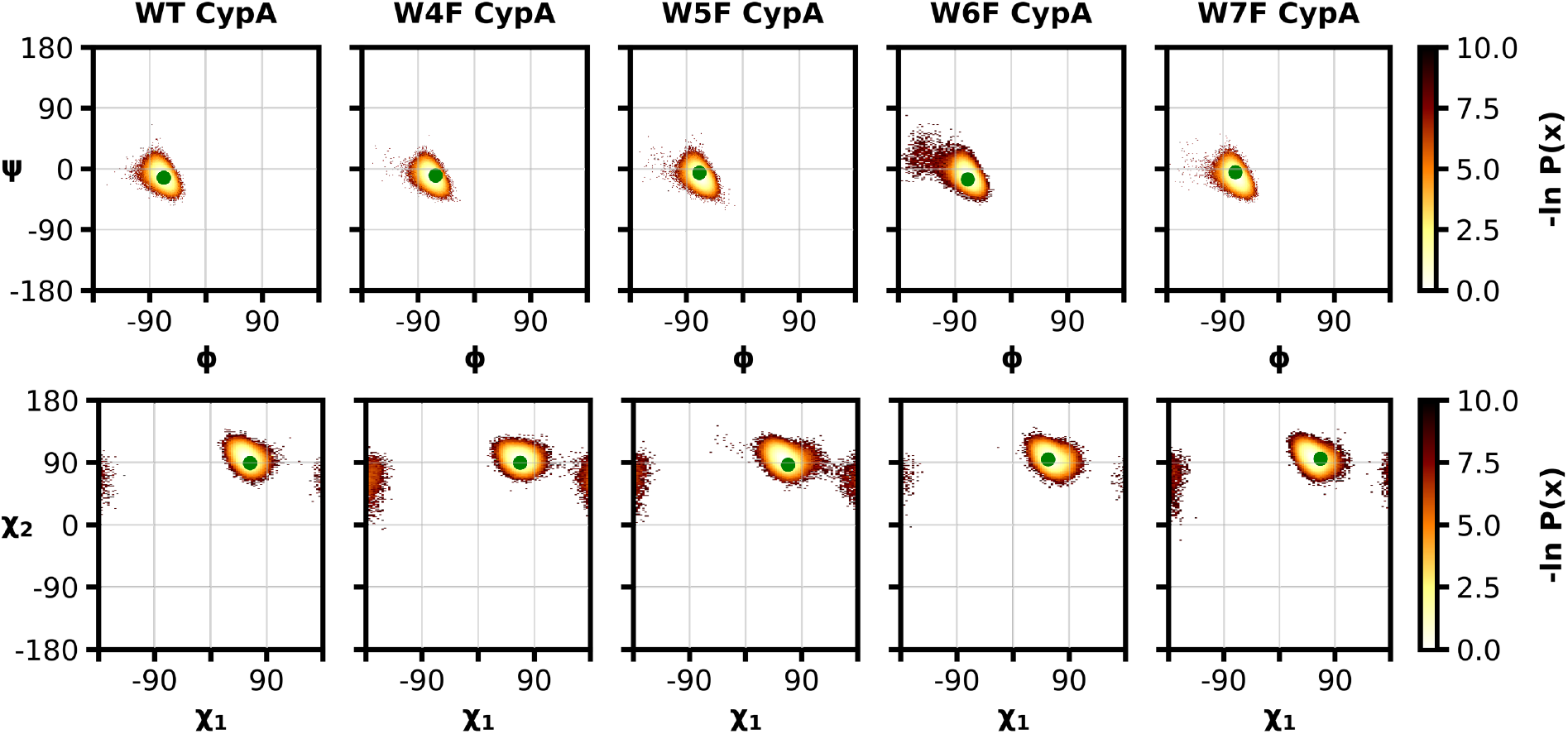
Ramachandran (Φ and Ψ) and Janin (χ_1_ and χ_2_) plots for an aggregate simulation time of 5 μs for fluorinated or native CypA proteins. The green filled circles indicate the original dihedral angle in either the native CypA crystal structure^63^, or the fluorinated CypA crystal structures at the respective 4, 5, 6, or 7 positions of the indole ring (unpublished data).

When evaluating the backbone RMSD and secondary structure predictions on a per-residue basis (**Figure S4**) excellent stability was maintained throughout multiple simulation replicates. However, a sharp increase in RMSD and corresponding change in secondary structure was noted between residues 146-153 and 102-107 during a fraction of both the wildtype and fluorinated CypA simulations. We isolated the characteristics of these increases in RMSD and found that they were largely driven by compensatory peptide-plane flips, which occur when changes in |Ψ_i_| + |Φ_i + 1_| are large, while changes in |Ψ_i_ + Φ_i + 1_| are comparatively small^82^. We accessed the per-residue level torsion angles of each trajectory (**Figure S5**) and found that residues 146-153 are a part of a loop which undergoes a hinge motion that’s compensated by a peptide-plane flip of residues 149 and 150 (top panel: **Figure S6**). Furthermore, residues 102-107 are a part of a beta turn that becomes twisted throughout our simulations and is compensated by peptide-plane flips of resides 103/104 and 107/108 (bottom and middle panel: **Figure S6**).

### C. ^19^F Relaxation of CypA

In addition to structural properties, we also evaluated whether longitudinal and transverse fluorine NMR relaxation rates would be accessible from our aggregate CypA simulation data. NMR relaxation rates function as useful probes to study dynamics and depend on the properties of a nucleus, modulated by its local environment. Fluorine relaxation is governed by both dipole-dipole interactions with neighboring proton spins as well as chemical shift anisotropy (CSA)^5^. Fluorine CSA in solution is averaged due to molecular tumbling, although CSA associated motions can cause oscillations of the local magnetic field that contribute to relaxation. Thus, fluorine relaxation is complex and not well understood.

In a first approximation, we calculated longitudinal and transverse relaxation rates from our aggregate CypA simulation data using an overall rotational correlation time τ_c_ of 8.2 ns^77^ (methods section G). We compared these calculated rates to the measured experimental rates^66^ (**Figure 7**). Overall, our calculated relaxation rates from aggregate MD simulation data are in good agreement with the experimental values. The average calculated longitudinal rates (R_1_) are systematically somewhat smaller than the experimental values, although within a reasonable error (**Table S4**). 4F-Trp CypA exhibited a somewhat larger R_1_ value (~ 2 s^−1^) than all others, which are similar and grouped around 1 s^−1^. The calculated transverse relaxation rates (R_2_) are not as close to the experimental ones, but still followed the same trend. They can be grouped into two sets comprising 5F- and 6F-Trp CypA with R_2_ values around 60-80 s^−1^, and 4F- and 7F-Trp CypA with R_2_ values around 110-120 s^−1^. Overall, the calculated relaxation values follow the same trends and agree well with the experimental ^19^F NMR relaxation rates.

**Figure 7.**
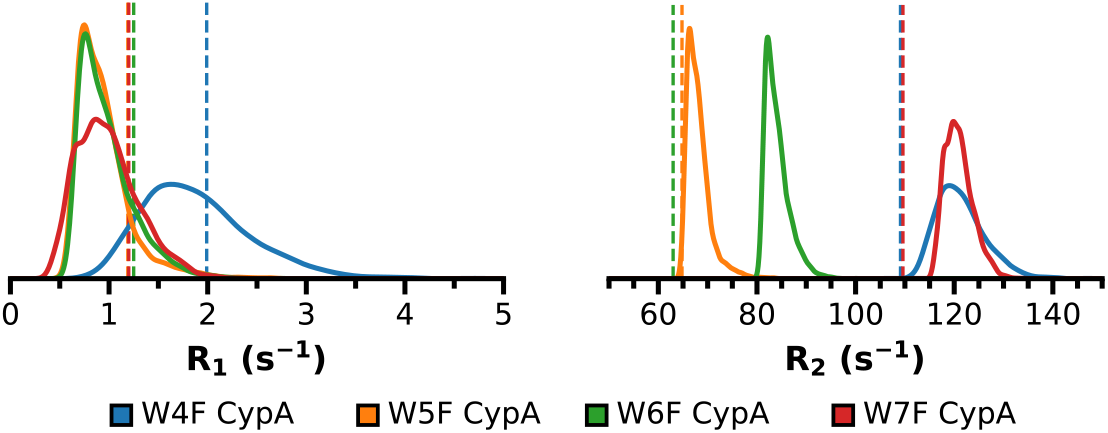
Fluorine longitudinal (R_1_) and transverse (R_2_) NMR relaxation rate distributions extracted from the final 80% of the 5 μs of aggregate MD simulation data for each fluorinated CypA variant. Experimental relaxation rates^66^ are shown as dotted vertical lines.

At this juncture it should be pointed out that our methodology for calculating relaxation rates from MD simulation data includes several key assumptions. In particular, our calculations assume that (i) the tryptophan sidechain essentially moves like the overall protein, i.e. it does not exhibit internal motions in addition to the overall molecular tumbling of the protein (ii) only hydrogens within a radius of 3 Å around the fluorine atom contribute to the relaxation rate, and (iii) any contribution from dipole-dipole and CSA cross-correlated relaxation^76^ is unaccounted for, since they may only contribute 10-25% to the total R_2_ relaxation rate^66^. Furthermore, the experimental relaxation rates are extracted by using single exponential fitting of the signal intensity decays, although multi-exponential fitting may more accurately capture crosscorrelation induced relaxation^66,74,83^. Despite these differences between simulation and experiment, as well as their respective assumptions, the simulation-based calculated rates agree surprisingly well with the experimental relaxation rates. In the future, potentially more accurate methods^84^ for fluorine relaxation rate calculations from MD simulations can be tested, which will help to dissect which components predominantly affect relaxation decay rates.

## V. Conclusions

We have reported the development and validation of force field parameters for a set of eight fluorinated amino acids that are commonly used for ^19^F NMR, for use with the AMBER ff15ipq protein force field. Our parameters include 181 implicitly polarized atomic charges and 9 unique bonded terms for 4, 5, 6, and 7 fluoro-tryptophan; 3, and 3, 5 fluoro-tyrosine; as well as 4 fluoro- and 4 trifluoromethyl-phenylalanine. We validated that our new parameters maintain the expected conformational propensities of the fluorinated amino acids consistent with both the respective canonical residues and previously characterized experimental X-ray structure-derived propensities extracted from structures deposited in the PDB. Fluorinated amino acid containing proteins, such as CypA, maintain the overall globular protein fold over multiple μs-timescale simulations and extracted ^19^F NMR relaxation rates are in good agreement with the corresponding experimental rates.

Overall, our results demonstrate the robustness of the “sweeping optimization” approach using the mdgx program of AMBERTools20 distribution^60^ and the power of the IPolQ lineage of AMBER force fields for modeling fluorinated proteins. Our workflow is readily applicable to other residue classes^32^ and can be easily expanded to include other fluorinated amino acids, if so desired. On the practical side, our force field parameters have numerous potential implications, particularly for use with complementary ^19^F NMR studies and when considering structural ensembles. Since ^19^F NMR data can be used to guide MD simulations and vice versa, our parameters will aid to derive an integrated, all-atom view of any fluorinated protein. Our parameters extend the macromolecular chemical space available to the AMBER ff15ipq force field and bridge the gap between computation and experiment for the collaborative study of fluorinated proteins at the atomic level.

Similar to the IPolQ methodology, the Force Balance approach and the corresponding AMBER-FB15 force field^85^ performs sweeping optimization of hundreds of parameters simultaneously, and includes nonlinear optimization methods which can incorporate alternative datasets, such as *in vitro* experiments, directly into the parameter optimization process. The FB15 force field has also recently been expanded to include parameters for phosphorylated amino acids^86^, but at this time, does not have parameters for halogenated residues such as the fluorinated amino acids presented in this work. In contrast to the IPolQ workflow, which is fully physics-based, the burgeoning development of machine learning based force fields^87^ are promising, but are heavily dependent on the training dataset being used. Thus, while physics-based approaches are more generalizable, deep learning methods can be more specialized for specific atoms or dependent on dataset properties. The goal of such machine learning based approaches is to narrow the gap between the efficiency of classical models and the accuracy of *ab initio* methods, by training a machine learning model to predict molecular potential energies, usually using quantum mechanical datasets. Examples of these approaches include the ANAKIN-ME^88,89^ neural network potential and the OrbNet^90^ method, which, once fully trained, can accurately predict the energetic properties of organic molecules within chemical accuracy of *ab initio* density functional theory calculations, but at a fraction of the computational cost. In the future, machine learning based methods may be readily paired with the IPolQ method to further enhance the efficiency of the parameter fitting and optimization process.

## Supporting Information

Supplemental figures include atom types for fluorinated amino acids and further analysis of CypA simulations: individual RMSD plots, carbonyl to backbone distances, secondary structure, per-residue RMSD, and per-residue backbone dihedral. Supplemental tables include individual force field bonded parameters, relative free energy differences between tetrapeptides, CSA tensors used for relaxation calculations, and comparison of average relaxation rates to experimental values.

## Acknowledgments

This work was supported by the NIH Pittsburgh Aids Research Training (PART) program grant T32AI065380 to D.T.Y., NIH grant P50AI150481 and NSF grant CHE-1708773 to A.M.G., and NIH grant R01GM115805 and NSF grant CHE-1807301 to L.T.C. Computational resources were provided by the University of Pittsburgh’s Center for Research Computing and through NSF XSEDE grant TG-BIO210161 to D.T.Y.

## Supplementary Figures

**Figure S1.**
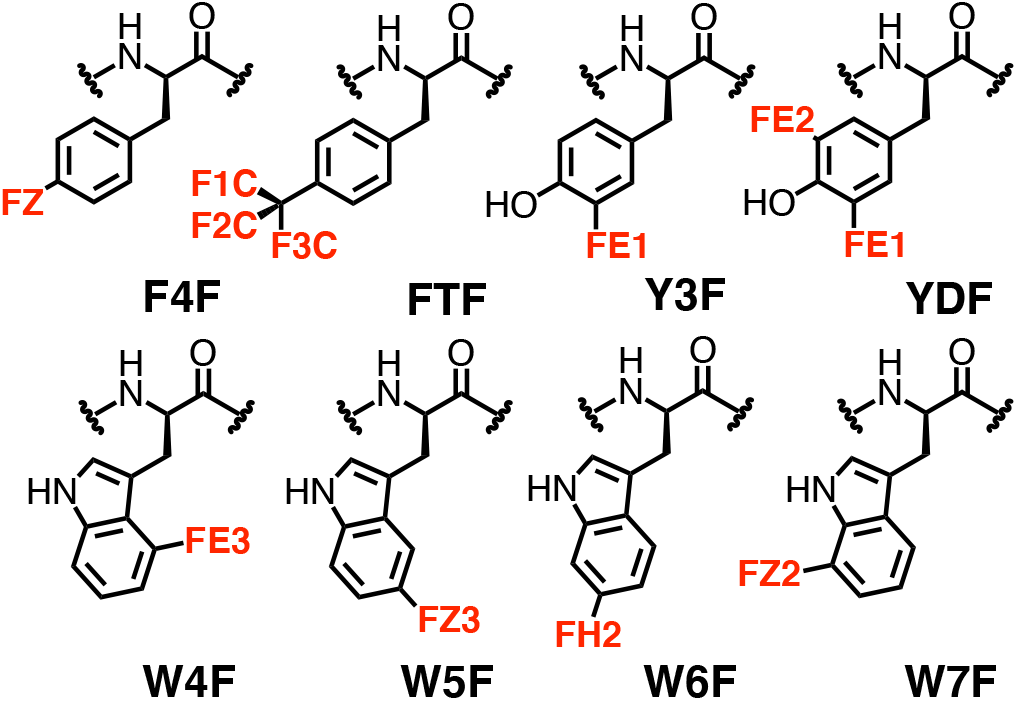
New fluorinated residue atom types presented in this work and their respective 3-letter identifier codes. The substituted fluorine atom-type is shown in red.

**Figure S2.**
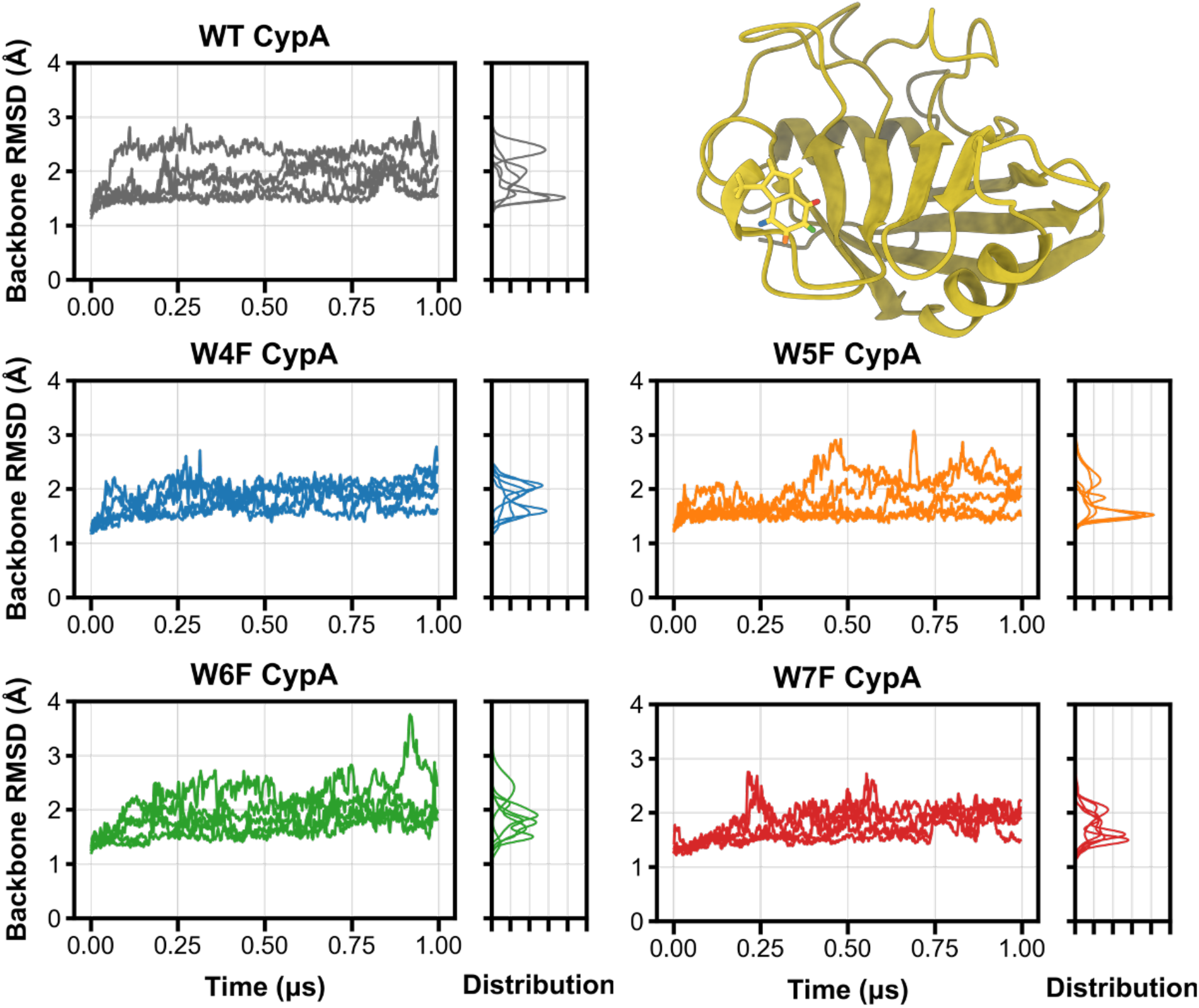
Backbone RMSD values with respect to the experimental X-ray structure (PDB: 3K0N)^63^ coordinates for wild-type and fluorinated CypA proteins extracted from each of the five independent 1 μs simulation replicates. The distribution of RMSD values is shown for each protein. The protein X-ray structure is shown in ribbon representation in the upper-right with individual fluorine positions on the Trp121 indole ring color coded like the trajectory plots.

**Figure S3.**
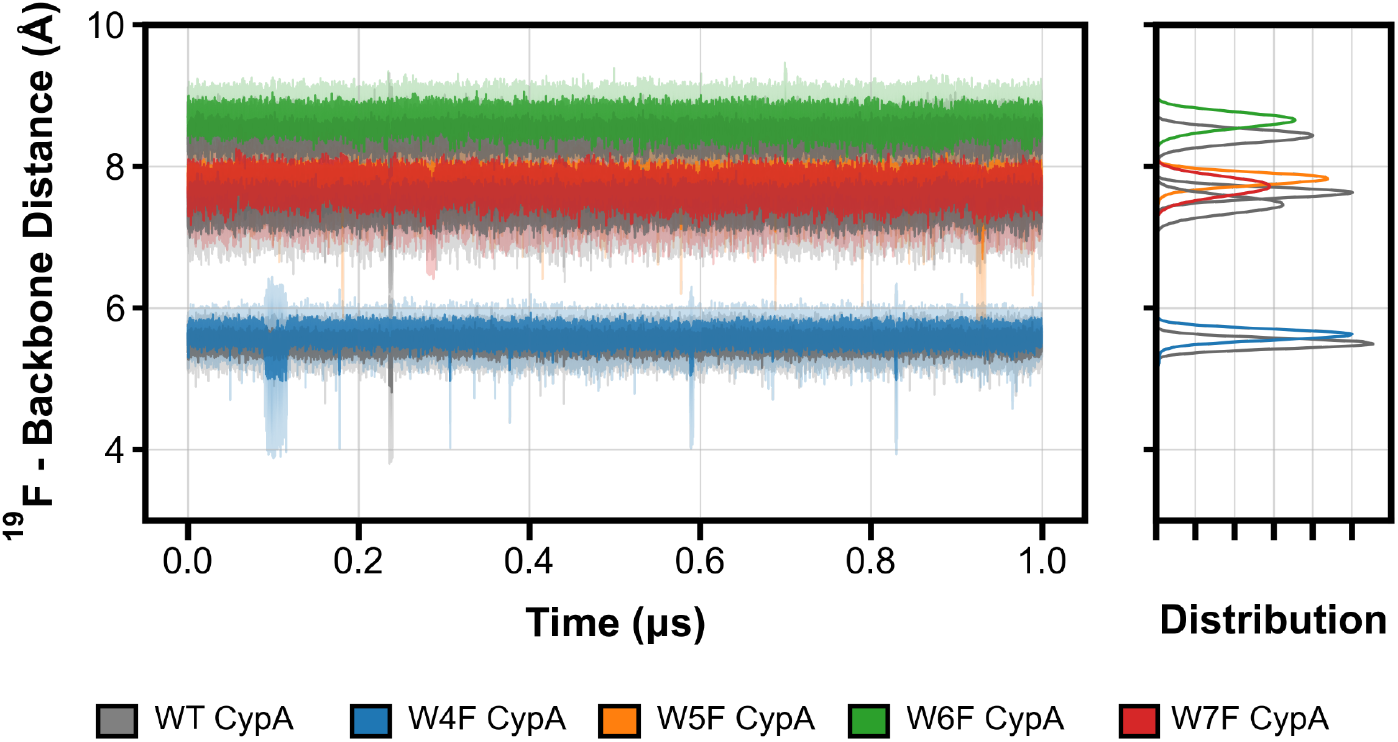
Average distance between the center of mass of the fluorine atom and the backbone carbonyl ± the standard deviation over the simulation time course for the fluorinated CypA variants. The distance for the hydrogen atom and the backbone carbonyl for wild-type CypA is shown in grey. This data represents all five independent replicate simulations of 1 μs each for each system.

**Figure S4.**
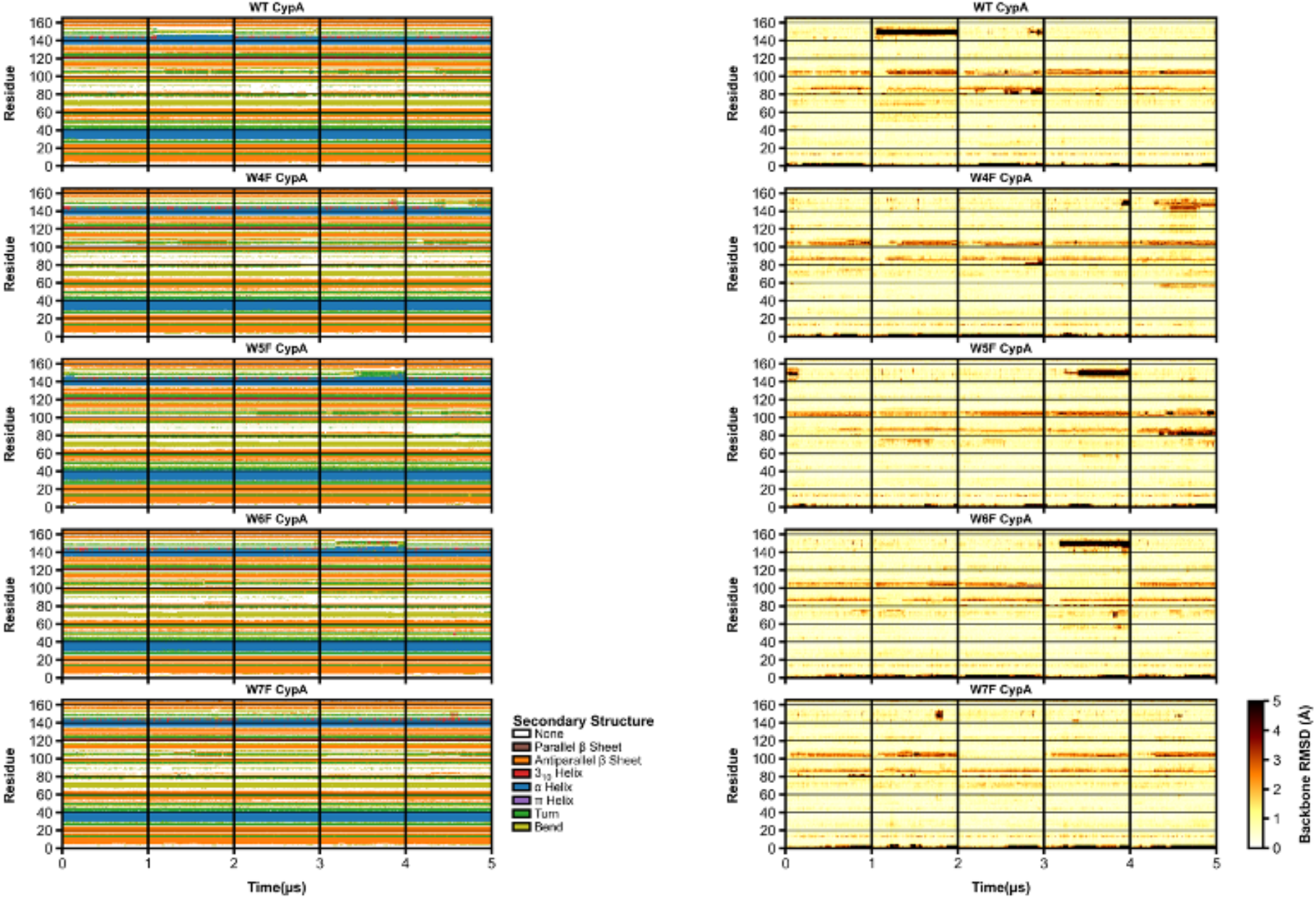
Secondary structure and per residue backbone RMSD values with respect to the crystal structure for both native CypA and the fluorinated variants. Each 1 μs simulation was replicated five times (separated by vertical lines) for a discontinuous simulation time of 5 μs for each system.

**Figure S5.**
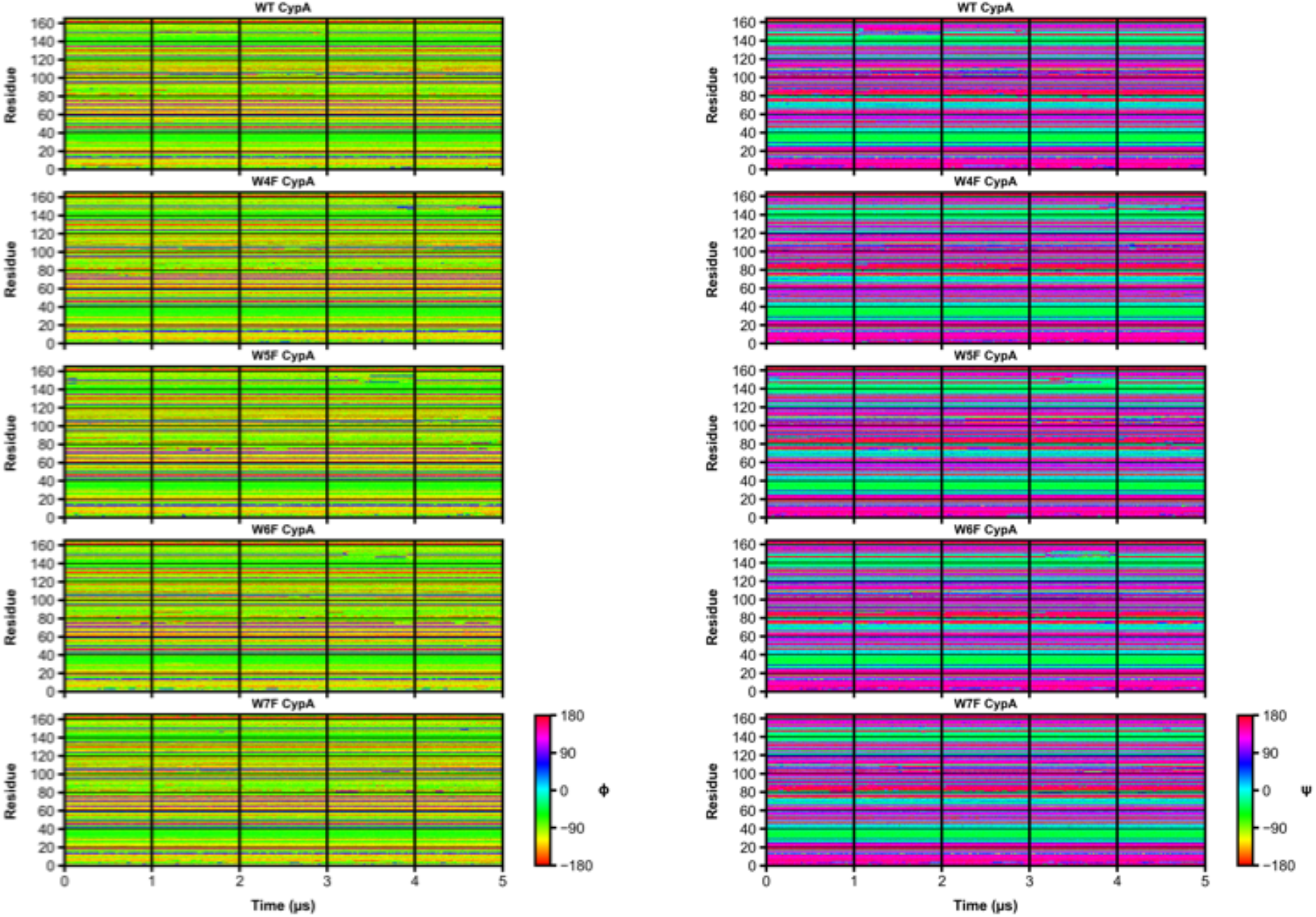
The per-residue Φ and Ψ dihedral angle variations for wild-type CypA and the fluorinated variants. Each 1 μs simulation was replicated five times (separated by vertical lines) for a discontinuous simulation time of 5 μs for each system.

**Figure S6.**
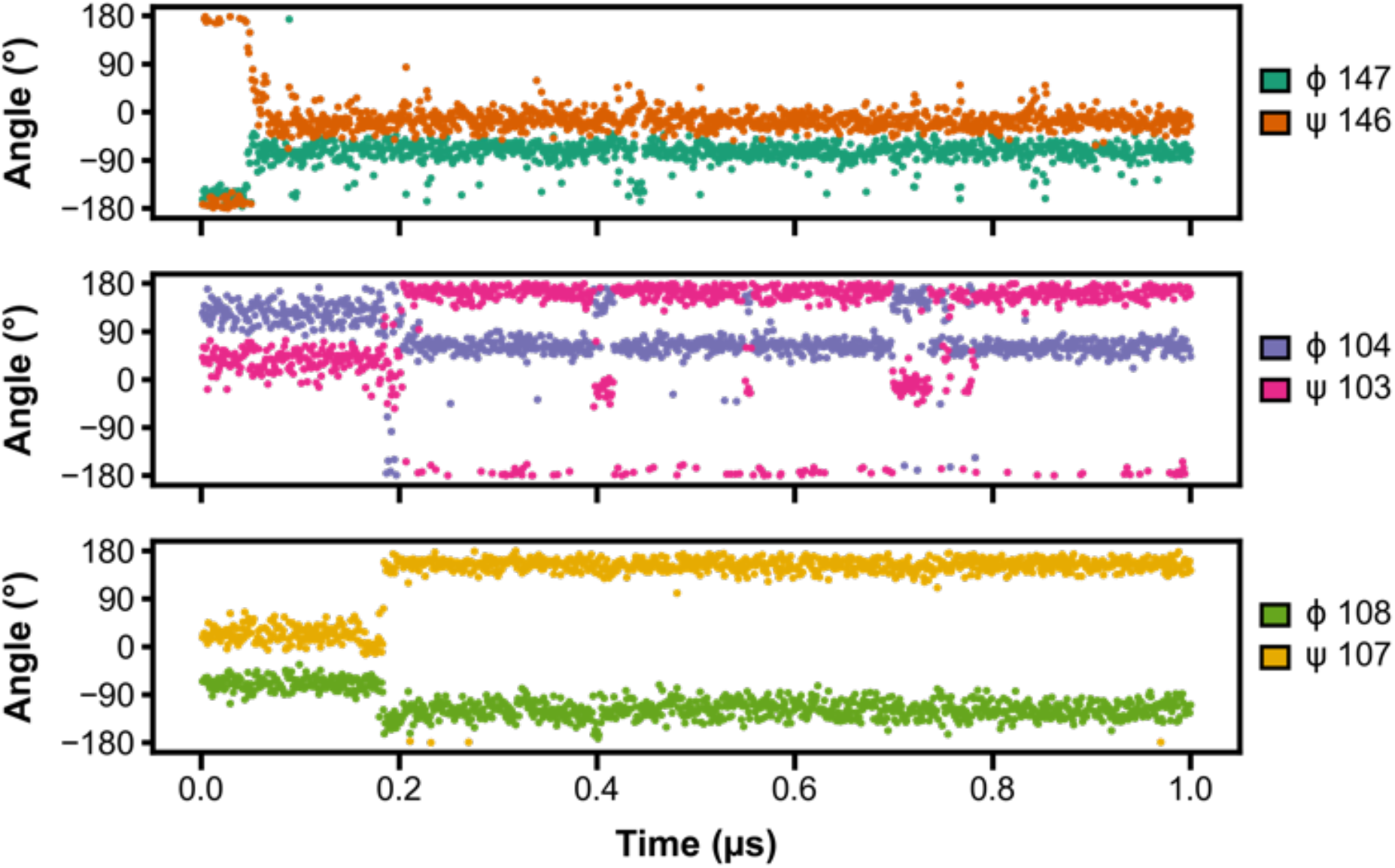
The Φ and Ψ dihedral angles of key neighboring residues around the amino acid exhibiting the jumps in backbone RMSD in **S4**, taken from a representative 1μs simulation of the non-fluorinated, wild-type CypA protein. Changes in adjacent dihedral angles are in opposite direction to the original jump, indicative of peptide-plane flips.

## Supplementary Tables

**Table S1.** Bond, angle, and torsion terms for the current set of fluorinated amino acid parameters.

| Bond | Force Constant<br>(kcal/mol/Å<sup>2</sup>) | Bond Length (Å) |  |
| --- | --- | --- | --- |
| 3C-CA | 317.4204 | 1.5100 |  |
| 3C-F | 367.0233 | 1.3800 |  |
| Angle | Force Constant<br>(kcal/mol/rad<sup>2</sup>) | Angle (°) |  |
| CB-CA-F | 109.3865 | 133.44 |  |
| CN-CA-F | 53.6158 | 118.76 |  |
| C—CA-F | 55.5980 | 120.15 |  |
| 3C-CA-CA | 70.9193 | 119.54 |  |
| CA-3C-F | 96.9340 | 111.90 |  |
| F—3C-F | 98.1332 | 107.08 |  |
| Dihedral | Barrier Height<br>(kcal/mol) | Phase Angle (°) | Periodicity |
| F-3C-CA-CA | 3.83580 | 0.0 | 2.0 |

**Table S2.** Average ± one standard deviation, minimum, and maximum ΔΔG values of the relative free energy difference of the tetrapeptide with a fluorine substituted residue, compared to the same peptide with a non-fluorinated amino acid.

| Peptide | Avg $\Delta\Delta G$ (kcal/mol) | Min $\Delta\Delta G$ (kcal/mol) | Max $\Delta\Delta G$ (kcal/mol) |
| --- | --- | --- | --- |
| W4F | 0.158 ± 0.162 | -0.312 | 0.637 |
| W5F | 0.089 ± 0.137 | -0.378 | 0.530 |
| W6F | -0.007 ± 0.139 | -0.626 | 0.547 |
| W7F | 0.062 ± 0.137 | -0.484 | 0.474 |
| Y3F | 0.088 ± 0.196 | -0.457 | 0.651 |
| YDF | 0.030 ± 0.228 | -0.554 | 0.551 |
| F4F | 0.273 ± 0.186 | -0.367 | 0.881 |
| FTF | 0.203 ± 0.268 | -0.534 | 0.859 |

**Table S3.** Experimental NMR principal components of the CSA tensor, reduced anisotropy, and asymmetry parameters for fluoro-substituted tryptophan powders^78^. These values were used to calculate the relaxation rates from MD simulations for each respective CypA variant.

| | $\sigma_{11}$ (ppm) | $\sigma_{22}$ (ppm) | $\sigma_{33}$ (ppm) | $\delta_\sigma$ (ppm) | $\eta$ |
| --- | --- | --- | --- | --- | --- |
| 4F-DL-Trp | 11.2 | -48.3 | -112.8 | 62.8 | 0.9 |
| 5F-DL-Trp | 4.8 | -60.5 | -86.1 | 52.1 | 0.5 |
| 6F-DL-Trp | 12.9 | -51.2 | -91.6 | 56.2 | 0.7 |
| 7F-DL-Trp | 4.6 | -48.3 | -123.3 | 67.6 | 0.8 |

**Table S4.** Comparison between the experimental ^19^F NMR relaxation rates and those extracted from MD simulations. Each value ± one standard deviation is listed.

| | Experimental | | Average $\pm \sigma$ from MD –<br>All protons < 3Å | |
| --- | --- | --- | --- | --- |
| System | $R^1$ (s<sup>-1</sup>) | $R^2$ (s<sup>-1</sup>) | $R^1$ (s<sup>-1</sup>) | $R^2$ (s<sup>-1</sup>) |
| W4F <u>CypA</u> | 1.99 ± 0.05 | 109.1 ± 4.9 | 1.89 ± 0.55 | 121.3 ± 4.94 |
| W5F <u>CypA</u> | 1.19 ± 0.02 | 64.8 ± 2.5 | 0.94 ± 0.27 | 68.1 ± 2.47 |
| W6F <u>CypA</u> | 1.25 ± 0.01 | 63.0 ± 3.9 | 0.97 ± 0.27 | 84.1 ± 2.47 |
| W7F <u>CypA</u> | 1.20 ± 0.01 | 109.6 ± 1.9 | 0.99 ± 0.31 | 120.9 ± 2.82 |

## Notes

### Competing Interest Statement

The authors have declared no competing interest.

